# Microtubule length correlates with spindle length in *C. elegans* meiosis

**DOI:** 10.1101/2023.11.23.568459

**Authors:** Vitaly Zimyanin, Stefanie Redemann

## Abstract

The accurate segregation of chromosomes during female meiosis relies on the precise assembly and function of the meiotic spindle, a dynamic structure primarily composed of microtubules. Despite the crucial role of microtubule dynamics in this process, the relationship between microtubule length and spindle size remains elusive. Leveraging *C. elegans* as a model system, we combined electron tomography and live imaging to investigate this correlation. Our analysis revealed significant changes in spindle length throughout meiosis, coupled with alterations in MT length. Surprisingly, while spindle size decreases during the initial stages of anaphase, the size of antiparallel microtubule overlap decreased as well. Detailed electron tomography shows a positive correlation between microtubule length and spindle size, indicating a role of microtubule length in determining spindle dimensions. Notably, microtubule numbers displayed no significant association with spindle length, highlighting the dominance of microtubule length regulation in spindle size determination. Depletion of the microtubule depolymerase KLP-7 led to elongated metaphase spindles with increased microtubule length, supporting the link between microtubule length and spindle size. These findings underscore the pivotal role of regulating microtubule dynamics, and thus microtubule length, in governing spindle rearrangements during meiotic division, shedding light on fundamental mechanisms dictating spindle architecture.

## Introduction

Female meiosis is a complex cellular process that involves two successive divisions, in which half of the chromosome content is extruded into two polar bodies, resulting in the formation of a haploid oocyte that is ready for fertilization. Errors in the segregation of chromosomes often lead to aneuploidy, an abnormal number of chromosomes, which has detrimental consequences for the development and survival of the oocyte. It is known that misregulation of microtubule (MT) dynamics disrupts meiosis (1,2). However, the mechanisms that explain this observation have remained opaque.

During the meiotic cell division, the segregation of chromosomes is mediated by the meiotic spindle, a self-assembling, dynamic structure composed of MTs, motor proteins, MT binding proteins, and various regulatory factors. Unlike other spindles in most animals, female meiotic spindles typically have no centrosomes, due to centriole elimination during oogenesis. Thus, female meiotic spindles must assemble and segregate chromosomes without MT organizing centers. Specifically, this is true in humans (3), mice (4) and also in nematodes (5). The assembly and regulation of meiotic spindles involves multiple stages, including MT nucleation, organization and stabilization, as well as the coordination of motor protein activity and regulatory signaling. While much is known about the role of motor proteins and regulatory signaling (6–9) in meiosis, comparatively less is known about the role and importance of MT dynamics during meiotic spindle formation and chromosome segregation (10), even though dynamic MTs are the main building blocks of spindles. Several publications showed that the microtubule severing protein katanin plays an important role in meiotic spindles by regulating microtubule length, turnover dynamics, and organization. Katanin’s ability to sever microtubules contributes to spindle remodeling, ensuring the proper assembly, function, and reorganization of the meiotic spindle during cell division (11–16). While it is known that MTs turn over at high rates in spindles, and that this can generate forces and modify the overall connectivity of the spindle (17), it has remained elusive how this process is regulated and how it contributes to essential spindle functions such as chromosome congression and chromosome segregation in anaphase.

Specifically for meiotic divisions previous research has shown that MT growth dynamics are crucial for correct meiotic spindle formation and function, as treatment of mouse oocytes with taxol impairs MT and chromosome organization as well as meiotic progression and leads to aneuploidy (1). Further, mutations or disruptions in proteins involved in MT de-/polymerization, such as XMAP215/Dis1 family proteins (18–22) and their regulators or MCAK (23–26) result in abnormal spindle structures, misaligned chromosomes, and failed chromosome segregation in yeast, Drosophila, mice and various other organisms. In addition, oocytes from older mice exhibit altered spindle MT dynamics which may contribute to age-related increase in chromosome segregation errors that are observed in these animals (2). These findings highlight the critical role of MT dynamics in establishing functional meiotic spindles and ensuring accurate chromosome segregation.

One of the main obstacles in identifying the role of MT dynamics during meiosis is the limitation of light microscopy in resolving submicron structures. Thus, information such as the number and length of MTs, their interactions with chromosomes, their nucleation pattern and conformation cannot be obtained by light microscopy and is only inferred from indirect measurements. To overcome these shortcomings, we are using 3D electron reconstructions of spindles in *C. elegans* (27–30). *C. elegans* has been extensively used as a model system to understand chromosome segregation in meiotic and mitotic spindles, especially since most of the proteins involved in meiotic divisions in *C. elegans* have orthologues in humans (31).

Our previously generated reconstructions of complete spindles in *C. elegans* oocytes in different stages of meiosis I and II by 3D electron tomography revealed that these spindles are mainly composed of an array of short MTs with nearly half of them being shorter than 500 nm (32). It also showed that there are significant changes in MT number and length throughout meiosis, with the number of MTs doubling from metaphase to early anaphase and then reducing again by 2-fold in mid-anaphase. This change in MT number is consistent with light microscopy observations that MTs are highly dynamic in meiosis, with turn-over rates of approximately ∼ 5s and a rapid poleward movement with a rate of 8.5 ± 2.2 μm/min (32). Overall, our data suggested that regulation of MT dynamics, such as nucleation rate, growth and catastrophe rates, leading to changes in MT numbers and length might be involved in transitioning through meiosis.

In this study we combine static, yet spatially resolved, ultrastructural data obtained by electron tomography with dynamic data from light microscopy data to further determine which MT parameters and changes in organization could be critical for the rearrangement of the meiotic spindle structure throughout meiosis in the *C. elegans* oocyte.

## Results

### The microtubule overlap in the meiotic spindle decreases throughout anaphase

Live imaging of meiotic spindles during both meiotic divisions shows significant changes in the structure and size of the spindle throughout meiosis (Fig. 1). At metaphase of meiosis I, the average size of the meiotic spindle is 3.4μm± 0.13μm (SEM, n=7), it shortens significantly throughout the first ∼80 s of anaphase (2.1μm± 0.2μm (n=7)) and then increases in size during chromosome segregation while further progressing through anaphase (3.35μm± 0.4μm (n=7) (Fig, 1A). The meiotic spindle undergoes similar changes in meiosis II, where the average spindle size decreases from 3.9μm± 0.36μm (SEM, n=6) in metaphase to 2μm± 0.16μm (n=6) in early stage of anaphase and 3.1μm± 0.23μm (n=6) in late anaphase (Fig. 1B). This data shows that the meiotic spindle undergoes significant changes in length throughout meiosis and raises the question of how these changes are regulated.

**Figure 1.**
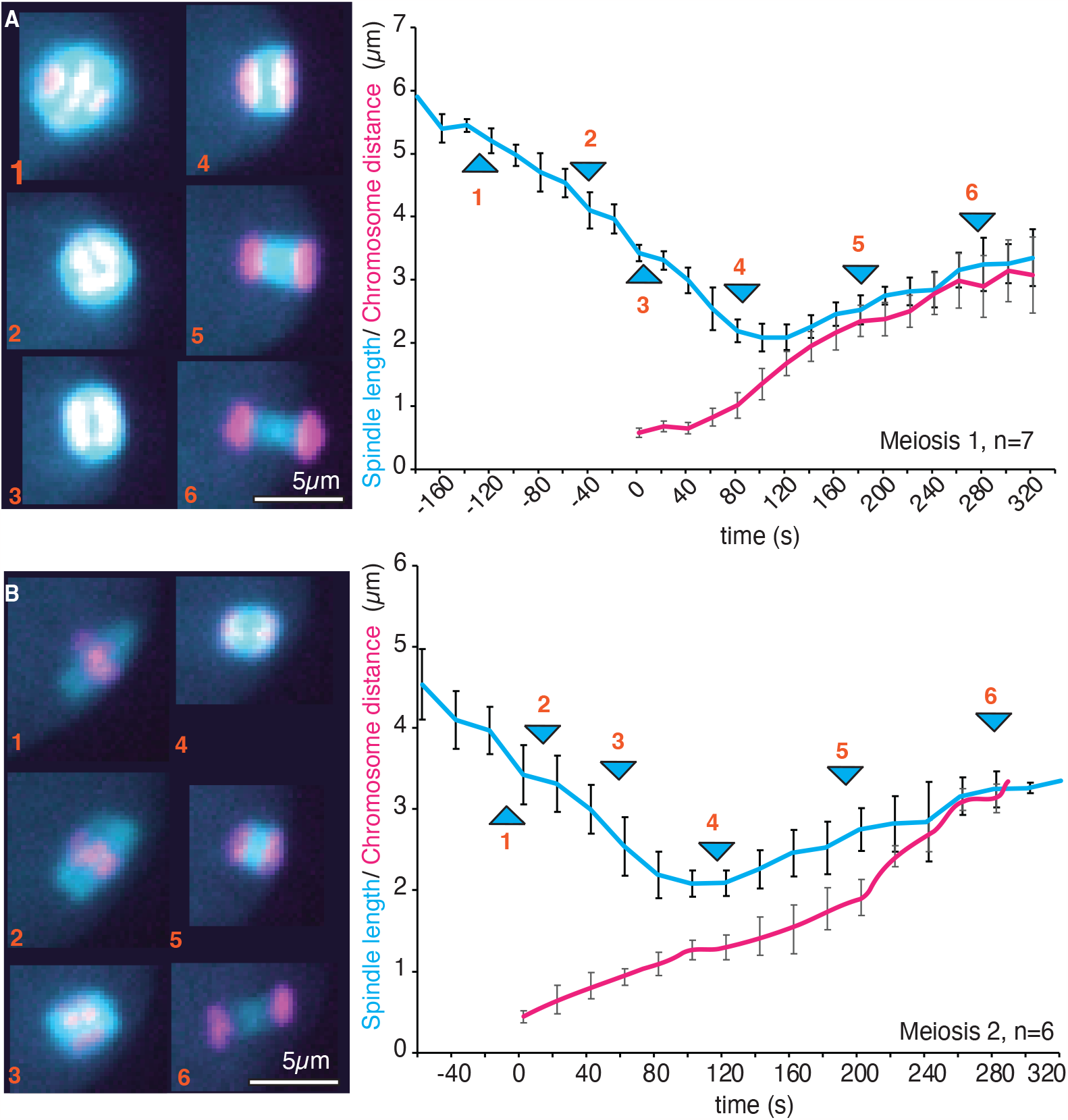
Meiotic divisions are accompanied by rearrangement of the spindle. **A**. Left panel. Fluorescent confocal images of different stages of *C. elegans* Meiosis I division labelled with GFP-Tubulin and Cherry-Histone. Right panel: Plot of the spindle length (blue) and chromosome distance (red) changes during Meiosis I. Arrows with numbers correspond to the stages of division shown in the images in the left panel. **B**. Left panel. Fluorescent confocal images of different stages of *C. elegans* Meiosis II Right panel: Plot of the spindle length (blue) and chromosome distance (red) changes during Meiosis II division. Arrows with numbers correspond to the stages of division shown in the images in the left panel.

One possible hypothesis is that motor proteins could drive an initial inward sliding of MTs in early stages of anaphase and thus lead to a decrease in spindle length. We speculate that such behavior could affect the region of antiparallel MT overlap in the spindle midzone, the region between the chromosomes during anaphase. The spindle midzone has been shown to actively participate in chromosome segregation and spindle elongation in mitosis and meiosis (33) presumably by generating outward directed forces. Our data on mitotic cells showed that the overlap between antiparallel MTs remained surprisingly stable (34) throughout anaphase suggesting that MTs within this overlap must either polymerize or be constantly rearranged to maintain the antiparallel overlap while the spindle midzone extends.

To obtain further insight into the changes of the midzone throughout meiosis that could regulate the changes in spindle length we quantified the MT arrangement in the spindle midzone. We used second harmonic generation (SHG) microscopy and two photon fluorescence (TP) microscopy simultaneously to measure the polarity of the collective microtubules throughout the spindle in vivo (35) (Fig. 2A). Surprisingly this data showed that the antiparallel overlap of MTs within the meiotic spindle became smaller from metaphase to anaphase, while the spindles shortened (Fig. 2B). In agreement with this we previously found that MTs in the spindle midzone in meiosis are sliding apart with similar velocities as the chromosomes (33). While the antiparallel MT region decreases throughout anaphase, we found that the MT overlap always extended for about 20% of the spindle length when the data was normalized. Although the decrease in MT overlap is consistent with the midzone playing an active role in chromosome segregation and the generation of outward directed forces, it cannot fully explain the observed decrease in spindle length throughout the initial stages of anaphase.

**Figure 2.**
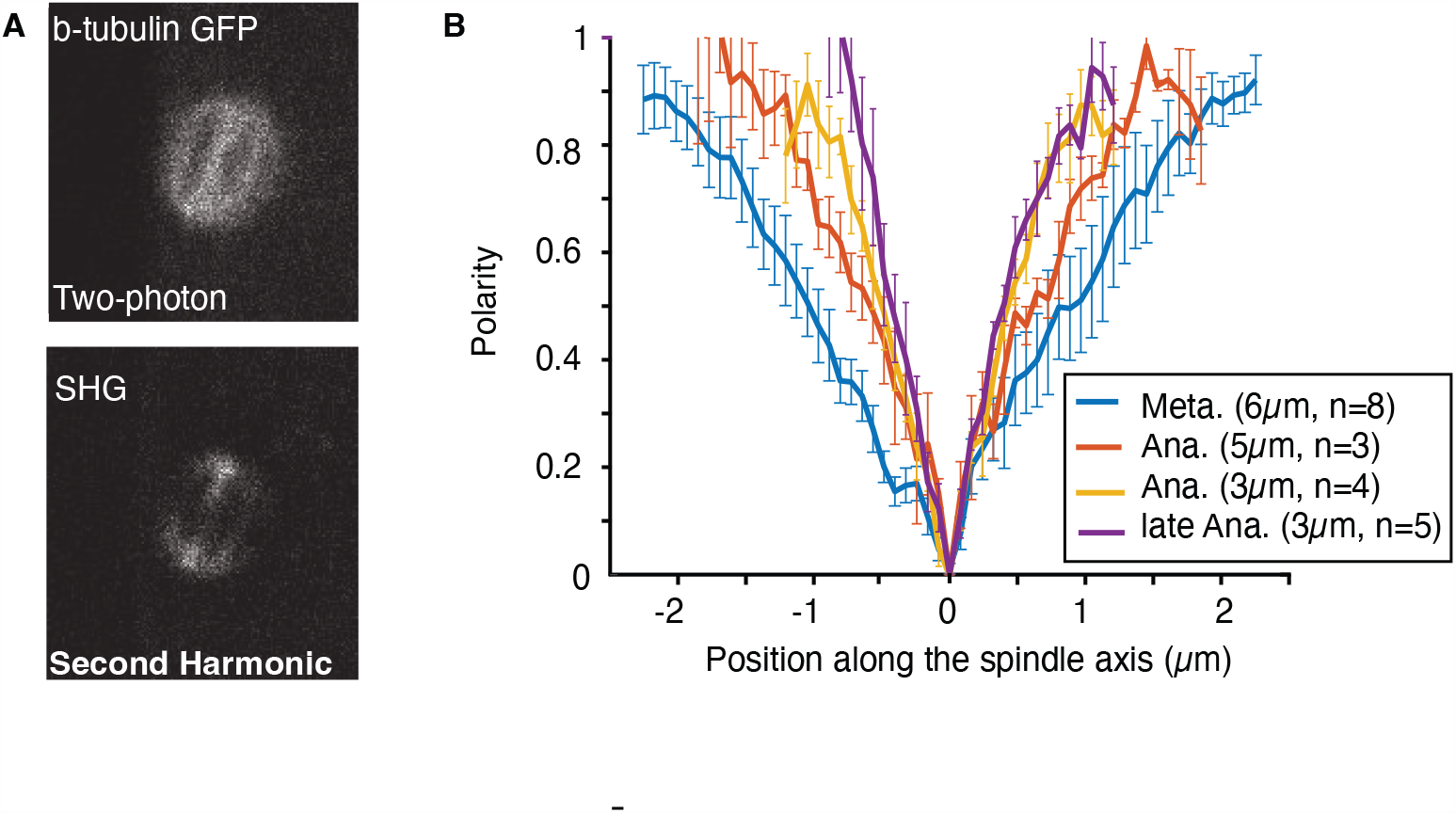
Microtubule overlap in the midzone is reduced during chromosome segregation. **A**. Two-photon fluorescence image (top) and second harmonic image (bottom) of a representative Meiosis I metaphase spindle showing overall microtubules localization (top) and their polarity (bottom): low signal indicating in SHG corresponding to antiparallel MT. **B**. Average polarity plots of microtubules in the spindle throughout metaphase and anaphase of meiosis I divisions obtained by the combination of SHG and TP microscopy. 0 on the x-axis represents the spindle center. Average spindle length and sample number is indicated in the legend. Error bars are sem.

### Microtubule length correlates with the spindle size throughout meiosis

To establish which characteristics of MTs change during the re-arrangement of meiotic spindles we used 3 D electron tomography to obtain detailed information about MT organization in meiotic spindles throughout meiosis I and II.

We previously reconstructed six meiosis I and four meiosis II (Fig. 3A,B) spindles at different stages throughout cell division (32,36). In agreement with the light microscopic data the reconstructed spindles show similar changes in spindle length throughout meiosis, with spindles shrinking from metaphase to early anaphase and then increasing in length while anaphase progresses (Fig. 3C top). These changes in size are accompanied by changes in MT number (Fig. 3C middle) and length (Fig. 3C bottom), where spindles in early anaphase that were presumably shortening showed increased MT numbers and decreased MT length (Fig.3C). To further determine changes in MT properties that could affect spindle length, we used the detailed data obtained by electron tomography to determine potential correlations between spindle length and MT number, average MT length, total polymer length (the added length of all MTs in the spindle) and chromosome volume. In total we re-visited 10 complete spindles with an average of ∼3500 MTs each. While this is a rather small sample size this data still provides valuable insights into potential correlations.

**Figure 3.**
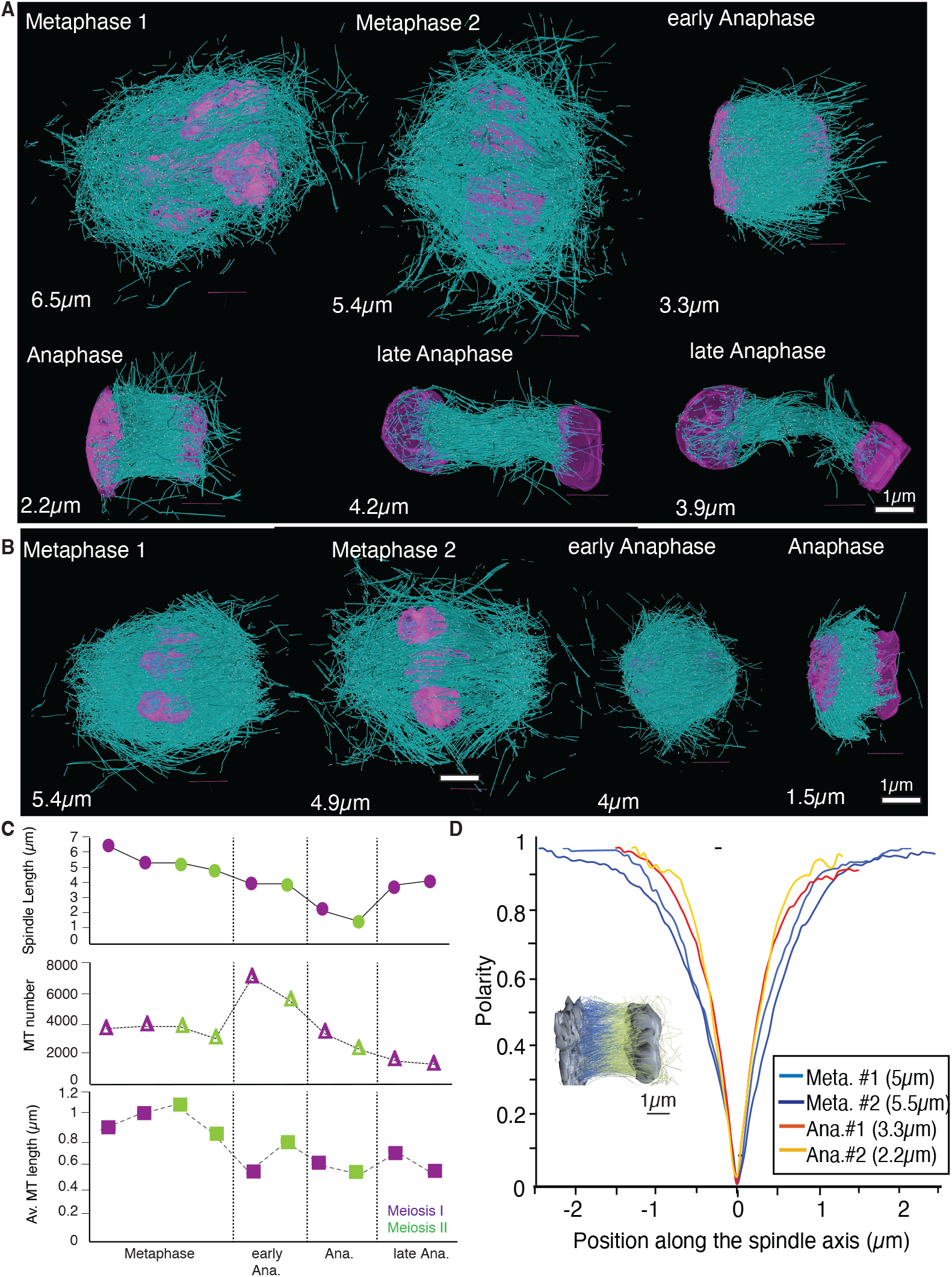
Ultrastructural TEM reconstructions provide structure of the meiotic spindle at the resolution of single MT. **A-B**. 3D tomographic reconstructions of different stages of Meiosis I (A) and Meiosis II (B) obtained by electron tomography. Scale bar is 1μm **C**. Plots showing different parameters of spindles determined by electron tomography in Meiosis I (purple) and Meiosis II (green) at stages shown in A-B, top: individual measurements of spindle length, middle: MT numbers of the individual spindles, bottom: Average MT length at the different stages **D**. Microtubule polarity plot based on the ultrastructural TEM reconstruction to show reduction of microtubule overlap with meiosis progression. 0 on the x-axis represents the spindle center. Average spindle length is indicated in the legend (n=1 for each spindle).

Quantification of the data suggest that the number of MTs (r=0.84, p<0.05) in a spindle as well as the average length of MTs (r=0.62, p<0.05) positively correlate with the total polymer length of MTs (Fig. 4 A,B,H). However, we could not show a correlation between MT length and MT number (r=0.12) (Fig. 4C,H). Our data also showed that there is no correlation between the MT number and the spindle length (r=0.07) (Fig. 4D,H), as well as between the total Chromosome Volume and the spindle length (r=-0.29, ns) in meiosis (Fig. 4E,H). Spindle length was only weakly and not significantly correlated with the total polymer length (r=0.46, ns) (Fig. 4G,H). Most strikingly our data revealed that there was a positive correlation between the average MT length and the spindle length (r=0.79, p<0.05) (Fig. 4 F,H) through meiosis. In summary this data suggests that the average length of MTs affects the spindle size and that MT dynamics are a more critical parameter in regulating spindle size. On the other hand, regulation of MT nucleation and thus MT numbers might play a less important role in meiosis.

**Figure 4.**
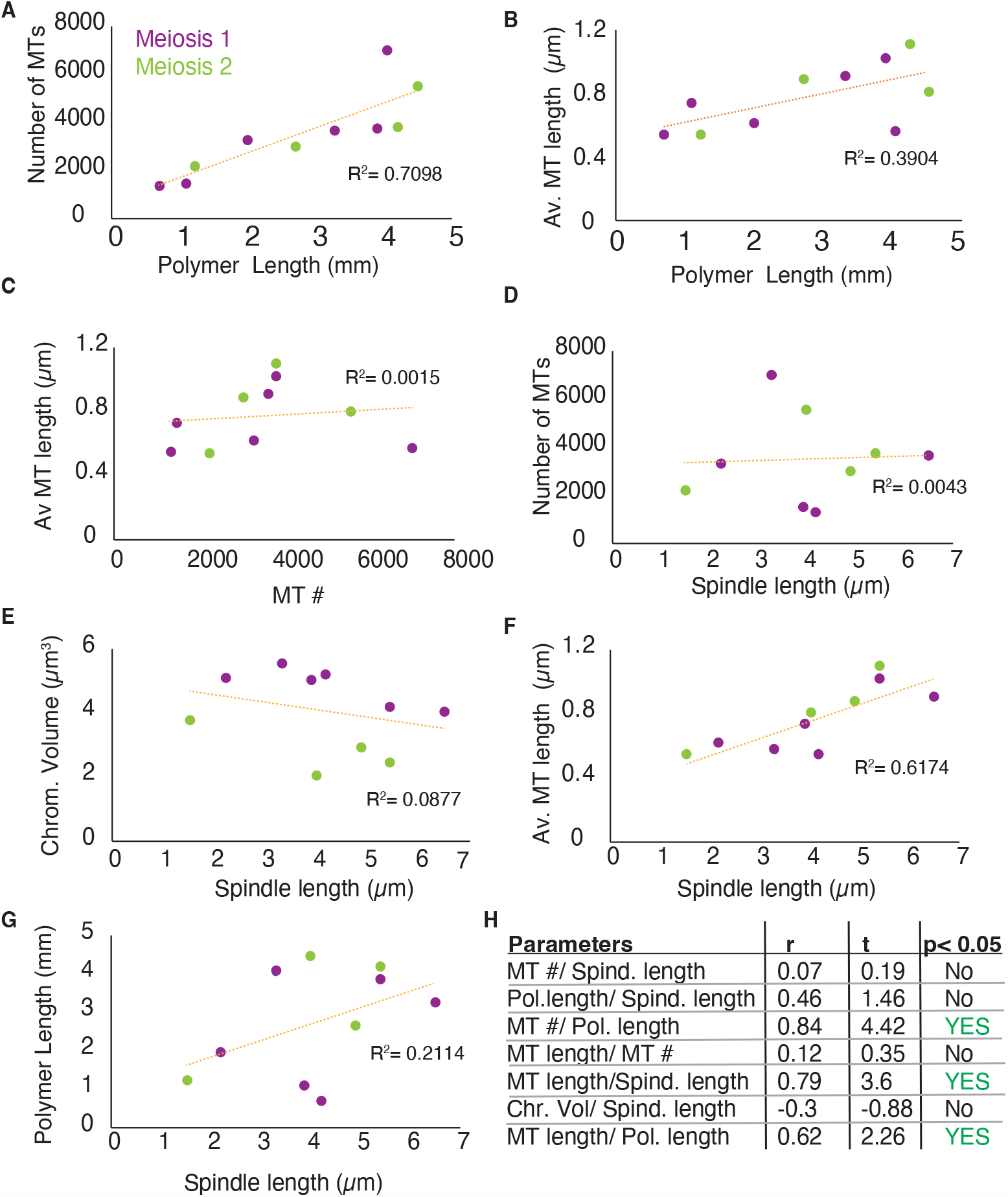
Microtubule length correlates with spindle length. **A-F**. Correlation plots demonstrating relationship between different parameters of spindles at different stages throughout Meiosis I (purple) and Meiosis II (green): Number of MTs, average MT length, Polymer length, Spindle length and Chromosome volume. **H**. Table showing correlation coefficient r, t and p-value significance for the analysis shown in **A-F**.

**Figure 5.**
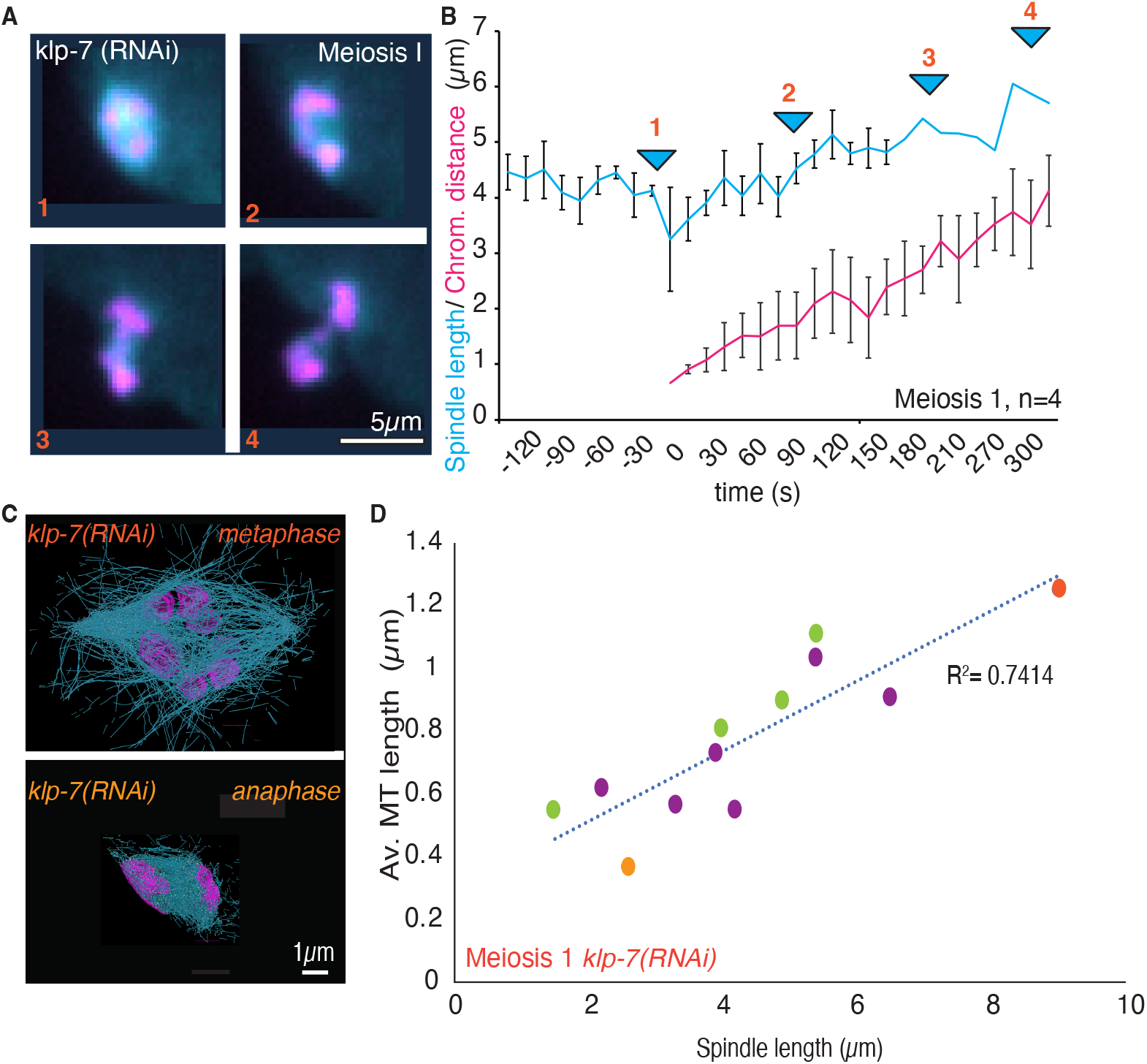
Microtubule and Meiotic spindle length are both increased in meiosis I metaphase spindles after *klp-7 (RNAi)*. **A**. Fluorescent confocal images of different stages of meiosis1 division in *klp-7 (RNAi)* treated embryos labelled with GFP-Tubulin and Cherry-Histone. **B**. Plot of the spindle length (blue) and chromosome distance (red) changes during meiosis I division. Arrows with numbers correspond to the stages of division shown in the images in the left panel. **C**. Reconstructions of microtubules in *klp-7 (RNAi)* meiotic spindles at metaphase (upper panel) and anaphase (lower panel) of the C. elegans one-cell embryo obtained by 3D electron tomography. **D**. Correlation plot comparing relationship of average microtubule length and spindle length in *klp-7 (RNAi)* spindles at metaphase (red) and anaphase (orange) shown in relation to the same values in wt meiosis I (purple) and II (green).

### Longer meiotic spindles are composed of longer microtubules

To further test the hypothesis that elongation of spindles corresponds to an overall increase in MT length, we quantified the effect of depletion of the MT depolymerase KLP-7. As a member of the kinesin-13 family, the *C. elegans* MCAK homolog KLP-7 possesses MT-depolymerizing activity, allowing it to selectively remove tubulin subunits from MT ends. This unique ability enables KLP-7 to influence MT length and stability. Analysis of meiotic spindles by light microscopy showed that KLP-7 depleted spindles do not display a shortening of the meiotic spindles during early anaphase in meiosis I, leading to longer spindles in early anaphase, with 2.1μm ± 0.18μm (n=7) in control vs 4.43μm± 0.36μm (n=4) after *klp-7 (RNAi)* at 80s after anaphase onset. We quantified two 3D tomographic reconstructions after *klp-7 (RNAi)*, one of a meiotic spindle in metaphase, which is elongated in comparison to control conditions, and one in anaphase, which is similar in size to control conditions. We found that in both stages the average MT length would be indicative of spindle length. Given the molecular function of KLP-7 this result supports our observations that changes in MT length is an important parameter and molecular mechanism that regulates spindle size in *C. elegans* meiosis.

## Discussion

Female meiosis is crucial for the formation of a viable oocyte and the precise chromosome segregation facilitated by the meiotic spindle is essential. Errors in this process often result in aneuploidy, underscoring the significance of understanding the underlying mechanisms governing meiotic spindle dynamics. Combining ultrastructural insights from electron tomography with dynamic observations from light microscopy, we aimed to identify MT parameters that might affect spindle morphology throughout the *C. elegans* oocyte division.

The meiotic spindle undergoes distinct changes in spindle size throughout meiosis, characterized by a notable shortening during early anaphase followed by elongation as anaphase progresses. This pattern is consistent across both meiotic divisions, underscoring the dynamic nature of spindle architecture throughout oocyte division. Assessing the MT overlap within the spindle midzone using second harmonic generation microscopy and two-photon fluorescence microscopy shows a reduction in antiparallel MT overlap from early anaphase throughout anaphase. This contrasts with our recently reported analysis of MT overlap in mitotic spindles, which remains constant throughout anaphase. However, our data on meiotic spindles shows that despite spindle size changes, the fraction of the antiparallel overlap region in comparison to the full spindle length remains relatively constant, implying some mechanism of overlap regulation during meiosis. Overall, this data suggests that alternative mechanisms might be at play regulating spindle length beyond changes in MT overlap.

Direct measurements of MT properties via electron tomography provide crucial insights into the potential correlation between MT properties and spindle length. Our analysis indicates a positive correlation between the length of MTs and the length of the spindles at different stages. Intriguingly, while MT length positively correlated with spindle size, there was no significant correlation between MT number and spindle length. This highlights the pivotal role of MT length in determining spindle size. This observation agrees with previous data that suggested that modulation of microtubule dynamics, in particular growth rates and catastrophe frequency, provides an efficient mechanism to control spindle length during embryonic development (37–39). It was also suggested that those changes in MT dynamics would affect the length of MT and that varying MT dynamics sets an average MT length that scales with cell volume and spindle length (39). Our data strongly supports the hypothesis that MT length affects spindle length, and that this correlation is not only functioning during development but also during the progression through meiosis.

Further reinforcing the link between spindle length and MT dynamics, the depletion of the MT depolymerase KLP-7 resulted in elongated spindles during early anaphase in meiosis I. Analysis of tomographic reconstructions showed an increase in MT average length and thus supported our observation, suggesting a strong association between MT length and resultant spindle size.

In conclusion, our integrative approach combining electron tomography with live imaging delineates the pivotal role of MT dynamics in governing spindle rearrangements throughout meiotic division. These findings underscore the importance of MT length regulation in determining spindle size and emphasize the need for further exploration to comprehensively understand the intricate interplay between MT dynamics and spindle architecture.

## Material and methods

### Worm strains gene silencing by RNAi

MAS91 (unc-119(ed3) III; ItIs37[pAA64; pie-1::mCherry::HIS58]; ruIs57[pie-1::GFP:: b-tubulin+unc-119(+)]) was used for experiments of fluorescence imaging, and fluorescence recovery after photobleaching. MAS91 was a gift from Martin Srayko. Worms were cultured and maintained at 16°C as described (40). RNAi experiments were performed by feeding (41). The feeding clone for *klp-7* was obtained from the RNAi library (42). L4 larvae were transferred to plates seeded with bacteria producing dsRNA and grown for 24 h at 25°C prior to analysis.

### Light microscopy

#### Sample preparation for light microscopy

Oocytes for live-cell imaging were prepared as described previously (43).

For FRAP measurements, meiotic spindles in oocytes were observed in the uterus of adult hermaphrodites (strain MAS91) mounted on a thin 4% Agarose pad between a slide and a coverslip. Polystyrene microspheres (Microspheres 0.10 μm, Polysciences, Inc.) were added to the agar solution before specimen mounting to immobilize the living worms.

#### Spinning disk confocal fluorescence imaging

Live imaging was performed using a spinning disk confocal microscope (Nikon Ti2000, Yokugawa CSU-X1), equipped with 488-nm and 561-nm diode lasers, an EMCCD camera (Hamamatsu), and a 60X water-immersion objective (CFI Plan Apo VC 60X WI, NA 1.2, Nikon). Acquisition parameters were controlled using a home-developed LabVIEW program (LabVIEW, National Instruments). Images were acquired every 1 second with a single z-plane.

### Sample preparation for electron tomography

Wild-type (N2) *C. elegans* embryos were collected in cellulose capillary tubes(44) and high-pressure frozen as described using an EM ICE high-pressure freezer (Leica Microsystems, Vienna, Austria)(45). Freeze substitution was performed over 2–3 d at – 90°C in anhydrous acetone containing 1% OsO_4_ and 0.1% uranyl acetate. Samples were embedded in Epon/Araldite and polymerized for 2 d at 60°C. Serial semi thick sections (200 nm) were cut using an Ultracut UCT Microtome (Leica Microsystems, Vienna, Austria) collected on Pioloform-coated copper slot grids and post stained with 2% uranyl acetate in 70% methanol followed by Reynold’s lead citrate. For dual-axis electron tomography (Mastronarde 1997), 15 nm colloidal gold particles (Sigma-Aldrich) were attached to both sides of semi-thick sections to serve as fiducial markers for subsequent image alignment. A series of tilted views were recorded using an F20 electron microscopy (Thermo-Fisher, formerly FEI) operating at 200 kV at magnifications ranging from 5000× to 6500× and recorded on a Gatan US4000 (4000 px × 4000 px) CCD or a Tietz TVIPS XF416 camera. Images were captured every 1.0° over a ±60° range.

### Quantification of electron tomography data

We used the IMOD software package (http://bio3d.colorado.edu/imod) for the calculation of electron tomograms(46). We applied the Amira software package for the segmentation and automatic tracing of microtubules(47). For this, we used an extension to the filament editor of the Amira visualization and data analysis software(29,48,49). We also used the Amira software to stitch the obtained 3D models in *z* to create full volumes of the recorded spindles(48,49). The automatic segmentation of the spindle microtubules was followed by a visual inspection of the traced microtubules within the tomograms. Correction of the individual microtubule tracings included manual tracing of undetected microtubules, connection of microtubules from section to section, and deletions of tracing artifacts (e.g., membranes of vesicles).

### Data analysis for tomography

Data analysis was performed using the Amira software (Visualization Sciences Group, Bordeaux, France).

#### Length distribution of microtubules

For the analysis of the microtubules length distributions, we previously found that removing MTs that leave the tomographic volume only had minor effects on the length distribution(48). Therefore, we quantified all microtubules contained in the volume. In addition, in all analyses, microtubules shorter than 100 nm were excluded to reduce errors due to the minimal tracing length.

#### Microtubule polarity

For this analysis the spindle was split into a left and a right half, then the position of each MT end was determined and MT were assigned a polarity according to the MT end position that was closest to the chromosomes/ spindle poles, with MTs that had an end closer to of the left side being colored in blue and MTs with ends closer to the right side in yellow. After this the number of MTs for each category (blue and yellow) along the spindle axis in 500nm steps was quantified and the ratio of blue to yellow was calculated. A ratio of 1:1 was converted to a polarity of 0. Based on this the polarity along the spindle length was plotted.

### Second harmonic generation imaging and two-photon florescence imaging

Simultaneous SHG imaging and TP imaging were constructed around an inverted microscope (Eclipse Ti, Nikon, Tokyo, Japan), with a Ti:sapphire pulsed laser (Mai-Tai, Spectra-Physics, Mountain View, CA) for excitation (850 nm wavelength, 80 MHz repetition rate, ∼70 fs pulse width), a commercial scanning system (DCS-120, Becker & Hickl, Berlin, Germany), and hybrid detectors (HPM-100-40, Becker & Hickl). The maximum scan rate of the DCS-120 is ∼2 frames/s for a 512 × 512 imge. The excitation laser was collimated by a telescope to avoid power loss at the XY galvanometric mirror scanner and to fill the back aperture of a water-immersion objective (CFI Apo 40× WI, NA 1.25, Nikon). A half-wave plate (AHWP05M-980) and a quarter-wave plate (AQWP05M-980) were used in combination to achieve circular polarization at the focal plane, resulting in equal SHG of all orientations of microtubules in the plane, unbiased by the global rotation of the spindle, the spatial variation in the angle of the microtubules, and the local angular disorder of microtubules. Forward-propagating SHG was collected through an oil-immersion condenser (1.4, Nikon) with a 425/30 nm filter (FF01-425/30-25, Semrock). Two-photon fluorescence was imaged with a non-descanned detection scheme with an emission filter appropriate for green-fluorescent-protein (GFP)-labeled tubulin in *C. elegans* (FF01-520/5-25, Semrock, Rochester, NY). Both pathways contained short-pass filters (FF01-650/SP-25, Semrock) to block the fundamental laser wavelength. Image analysis was performed with MATLAB (The MathWorks, Natick MA), and ImageJ (National Institutes of Health, Bethesda, MD.

### Statistical analysis

The number of measured samples is indicated in the main text or figure legends, and for all measurements experiments were performed on a minimum of three or more different days. Unless otherwise stated statistical analysis was performed by using Student *t*-test assuming equal variance and two-tailed distribution. The data were analyzed and tested using Prism, Excel or MATLAB software. Correlation was tested using the Pearson’s Correlation Coefficient.

Statistics are presented as mean ± SEM, The significance level was set at *p* < 0.05, and displayed on figures with asterisks: * *p* < 0.05, ** *p* < 0.01, *** *p* < 0.001, **** *p* < 0.0001.

## Acknowledgements

We are extremely grateful to Drs. Daniel Needleman and Che-Hang Yu for providing support and tools, as well as help with image analysis for the SHG and TP approach. The C. elegans strains were provided by the Caenorhabditis Genetics Center (CGC), which is funded by National Institutes of Health Office of Research Infrastructure Programs (P40 OD010440; University of Minnesota).

The Redemann lab was supported by 1R01GM144668-01 NIH NIGMS.

## Authors Contribution

S.R. generated and analyzed the light- and electron microscopy data, V.Z. helped with conceptualization and co-wrote the paper.

